# Statistical Analysis of Zika Virus Correlation to Microcephaly

**DOI:** 10.1101/046896

**Authors:** Pedro P. Simoni, Isabela C. Simoni

**Author notes:** Electronic address (Pedro P. Simoni).

## Abstract

A Statistical Analysis was performed to probe the correlation between the Zika Virus (ZIKV) and its possibility to induce Microcephaly in infants. It was found that without considering a false positive on tests for ZIKV on mothers there seems to be a statistical significance on ZIKV to cause Microcephaly in infants with a probability of 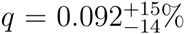. It was also shown that without knowing the confidence of the false positive tests of ZIKV it is not possible to statistically assert that this significance is true. It is proposed that to be able to discard the hypothesis of ZIKV not to cause the disease it is necessary to have a 30% of confidence in the ZIKV test.

## I. INTRODUCTION

The first isolation of ZIKV was made in 1947 from rhesus monkey in Zika Forest in Africa [1]. Since 2007 started to spread outside of Africa, Asia and the Pacific Islands. ZIKV major outbreaks have been occurring in the Americas since May 2015 with estimative of infecting more than 1.5 million people in Brazil and 50 000 around the world [2, 3]. ZIKV is a small enveloped virus belongs to the genus Flavivirus that contains a positive, single-stranded genomic RNA. In Brazil the mosquitoes Aedes aegypti is considered to be the main vector of the virus [4].It was is closely related to other flaviviruses of public health relevance including dengue, yellow fever and West Nile viruses [5].

In spite of its symptoms be mild, it seems that the ZIKV can cause central nervous system malformation such as Microcephaly in infants when their mothers get into contact with the virus in early stages of pregnancy [6]. To check this hypothesis it is proposed a statistical test to evaluate the confidence of this hypothesis (*H*_1_) over the Null Hypothesis (*H*_0_) of the ZIKV not to cause the microcephaly.
This paper is divided in the following manner: In Section II it is presented the statistical Methodology used to evaluate the scenario. Section III illustrates the main results on the statistical significance without considering false positive tests using only the statistical test of [7]. The problem is revisited in Section IV by considering the significance now adding a false positive assumption. Section V the analysis is done using the full data of [8] and Section VI presents a discussion on the interpretation of those results

## II. STATISTICAL METHODOLOGY

The first part of the analysis presented was based on the results of Schuler-Faccini, 2016 [7] where they collected 37 infants with microcephaly and obtained the result of 26 mothers (74%) reported symptoms related to ZIKV infections. To evaluate whether this result can be explained solely by the chance or if it is possible to probe the existence of a correlation between ZIKV and microcephaly it is necessary to take into consideration that the analysis on newborns was made by collecting babies known to suffer from microcephaly, so the fundamental question is what is the probability that given the babies have microcephaly, the mothers have ZIKV as well. Thus by probability theory,

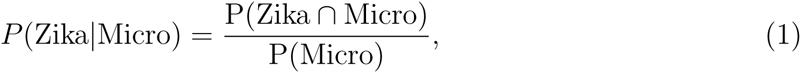

where *P*(Zika|Micro) is the probability to the mother is/was contaminated with ZIKV given that the baby has Microcephaly, *P*(Micro) is the total probability of developing microcephaly.

To obtain a meaningful result one must include four possibilities over the results obtained: (i) the mother may have the ZIKV and her baby microcephaly, but both are not correlated. (ii) The mother may have the ZIKV and her baby microcephaly, and ZIKV caused it. (iii) The baby has microcephaly but the mother does not have ZIKV virus and is a false positive and (iv) The baby does have a microcephaly and ZIKV at the same time, but the test does not found the ZIKV, i.e. a false negative.

i. The analysis of Schuler-Faccini, 2016 [7] was done with mothers from Brazil where it is known that microcephaly has a probability incidence of 0.5 cases in 10000, or a probability of *P*_*m*_ = 0.5 × 10^−4^ before the ZIKV outbreak. Also, the outbreak of ZIKV in Brazil has contaminated over 1.5 million of people in a country with 200 million people, thus the probability of the each mother to have ZIKV is *P*_*Z*_ = 3/600. Thus the probability of the mother to have both microcephaly and ZIKV at the same time with uncorrelated causes are *P*_*Z*_*P*_*m*_.
ii. Lets now suppose that the ZIKV has a probability q of in fact cause microcephaly, in this case the probability of a mother to have a disease and the baby microcephaly is *qP*_*Z*_. The case where there is no correlation is *q* = 0. Thus the conditional probability is,

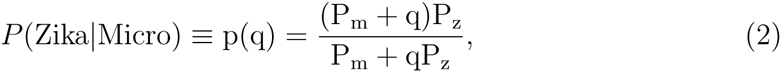

This means that *p*(*q*) is the function that describes the probability that a given person appears to have Zika virus in a test with only Microcephaly infants
iii. There is a probability f that the detected ZIKV infection is a false positive, that is, the mother seems to have ZIKV but does not. So that the probability of not having ZIKV but appears as a positive in the test is,
[1 — *p(q)]f*
It is not clear in the analysis of Schuler-Faccini, 2016 [7] what the confidence in the ZIKV infection is as it was only observed that the infants’ mothers presented symptoms compatible with ZIKV (which is similar to Dengue disease), so at first it is assumed *f* = 0 and check if there is a correlation and then to include the false positive test.
iv. There is a probability g that the detected ZIKV infection is a false negative, that is, the mother seems not to have ZIKV, but in fact she had/has it. Again the reliability of the test is not documented so at first it is assumed *g* = 0 and to check a possible correlation and then to include the false negative in the test. Therefore, the probability of being a false negative in the test is,
*p(q)g*
Thus, when considering both false negative and positive one should make the change,

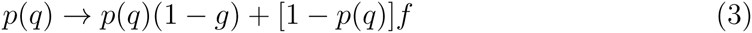

Table I summarizes all the input values for both analysis.

**TABLE I:** Parameters of the analysis.

| Parameter | Value | Meaning |
| --- | --- | --- |
| $P_Z$ | 3/600 | Probability of developing ZIKV disease |
| $P_M$ | $0.5 \times 10^{-4}$ | Probability of developing Uncorrelated Microcephaly |
| $q$ | Free Param. | Probability of ZIKV cause Microcephaly |
| $f$ | Free Param. | Probability of Detect ZIKV as a False Positive |
| $g$ | Free Param. | Probability of Not Detect ZIKV as a False Negative |

## III. ZIKA VIRUS AND MICROCEPHALY SIGNIFICANCE

It is possible to assume that all tests performed in this scenarios follows a binomial distribution that describes n uncorrelated tests with yes/no positives with k positives and a probability of having a positive to be *p*(*q*). This means that the probability of obtain k positive results in a n test is given by,

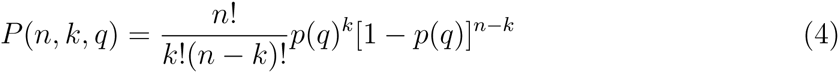

In [7], *k* = 26 and *n* = 37. Figure 1 shows the probability of obtain this result as a function of *q*, notice that the case where there is no relation between ZIKV and Microcephaly is for *q* = 0.

**FIG. 1:**
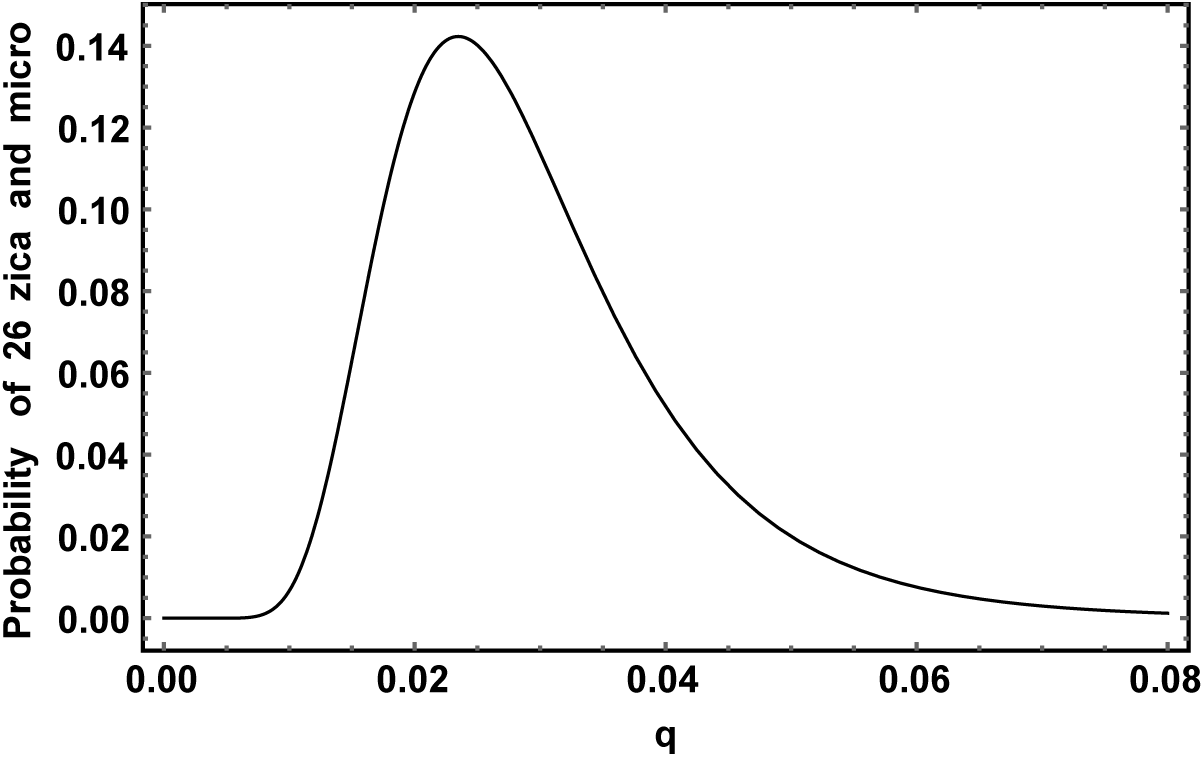
Probability of obtain 26 positives in a *n* = 37 sample of ZIKV and Microcephaly correlation study.

The probability has a clear peak *q* = 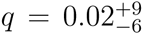 and rapidly fall to zero for *q* → 0. This suggests a strong correlation between the ZIKV and Microcephaly. This can be easily notice in the P-value test graphic in Figure 2 also, Table II summarizes all results obtained.

**TABLE II:**

| Variable | Analysis I<br>(Data: [7] ) | Analysis II<br>(Data: [7] ) | Analysis III<br>(Data: [8] ) | Analysis IV<br>(Data: [8] ) | Best Fit |
| --- | --- | --- | --- | --- | --- |
| $q$ | $2.3\%^{+15\%}_{-14\%}$ | 0 not Excluded | $0.092^{+15\%}_{-14\%}$ | 0 not Excluded | 0.12% |
| $f$ | 0 (fixed) | 60% (enough) | 0 (fixed) | 6% (enough) | 0.6% |
| $g$ | 0 (fixed) | $f = g$ | 0 (fixed) | 30% (enough) | 84% |

**FIG. 2:**
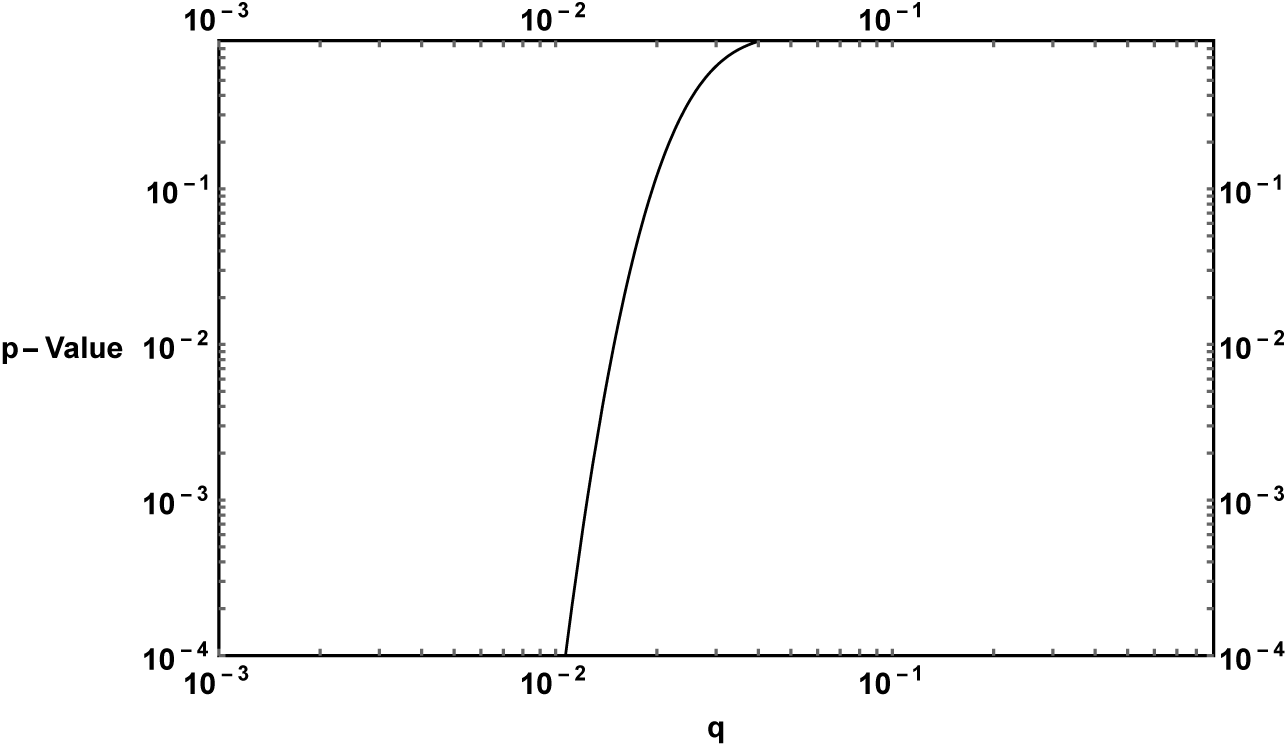
P-Value as a function of *q*.

It must be clear that Figure 1 is not a probability distribution function, it illustrate the probability of obtaining *k* = 26 positive tests given the probability *p(q)*. In contrast to *P(n, k, q)* which is indeed a probability distribution for the variable *k*.

## IV. CONSIDERATIONS ON FALSE POSITIVE TEST

Results of Section III may seem to indicate that indeed the ZIKV and Microcephaly are correlated and that microcephaly appears in 2.3% of the cases. But this analysis did not take into account the false positive and negative signal f and *g*. The *k* = 26 positive cases of study taken may have been contaminated by false positive yes and the remaining 11 cases not presenting ZIKV could be not detected false negatives. To take it into account both degrees of freedom it is necessary to modify Eq. (1) by taking *p(q)* → *p(q)*(1 — *g*) + *f*[1 — *p(q)*]. Unfortunately, both g and f are not well measured thus as a illustrative porpoise this section assumes *g* = *f*. In this case,

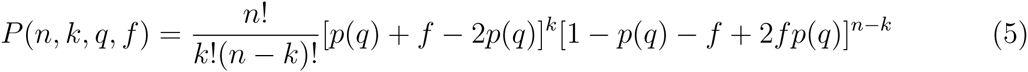

Now Figure 3 shows the new probability as a function of q assuming *f* = *αq* where represents the fraction of *q* that is in fact positive. The case where all the cases are false positives is *f* = 1 or *α* = 1/*q*. Notice that if f is big enough it is not possible to tell if there is correlation or not.

**FIG. 3:**
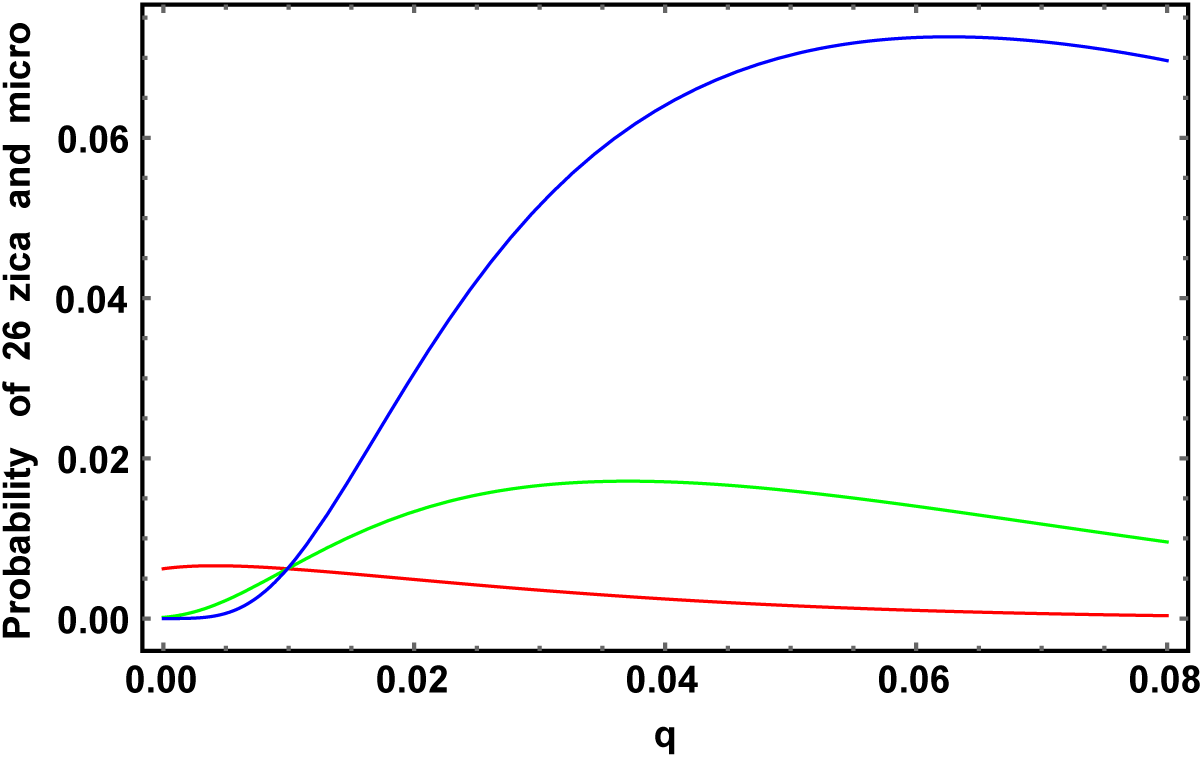
Probability of obtain 26 positives in a *n* = 37 sample of ZIKV and Microcephaly correlation study considering now the false positive and negative with *f* = *g* = *αq* where the red curve is for *α* = *0.5*, green if *α* = *0.3* and blue for *α* = 0.1.

The main question that arises is what is the optimal value for f so it can be distinguishable whether there is correlation or not between ZIKV and Microcephaly. Figure 4 helps to visualize this question, there it is plotted the P-Value of each assumption as a function of f. If its value is below the 0.05 threshold, there is a strong evidence against the hypothesis. In Figure 4 one can see the P-Value for both cases analyzed here, *q* = 0 and *q* = 0.023.

**FIG. 4:**
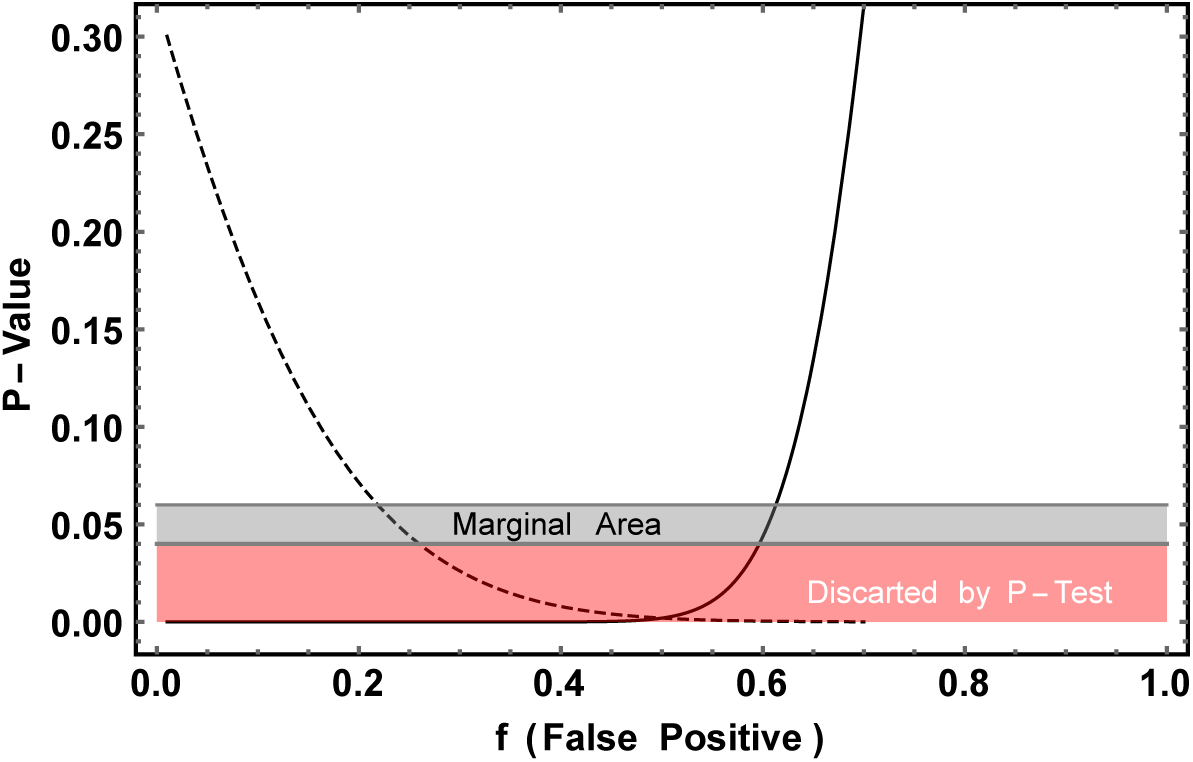
P-Value as a function of the false positive f. Full Line q=0 and dashed line *q* = 0.023.

Notice that the P-Value goes down for *q* = 0.023 as *f* increases in contrast it decreases for *q* = 0, this is a reflex of the fact that f is a measure of our ignorance, where *f* = 0 is no ignorance at all and *f* = 1 is total ignorance. With this graphic one can see that it is needed a false positive test bellow 0.4 or a 60% of hit of ZIKV test. This shows that even with small data points it would be possible to distinguish the two hypothesis if it was well known *f* and *g*.

## V. ECDC DATA ANALYSIS

The most recent results on the number of ZIKV and Microcephaly are presented in the ECDC website [8], there one can see that 462 cases of Microcephaly were studied resulting in 41 cases of Microcephaly and ZIKV at the same time. In spite of the great number of case study, it compiles several number of studies, each of them with its own *f* and *g* probabilities.

To take everything into account it is necessary to calculate the distribution function,

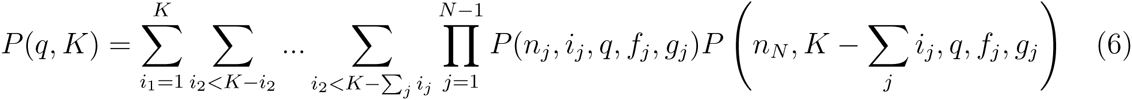

Where *N* is the number of experiments and *n*_*i*_,*f*_*i*_ and *g*_*i*_ are the parameters of each experiment and the sums should be done only for *i*_*j*_ < *n*_*j*_ and *K* = Σ_j_ *k*_*j*_. As *f*_*i*_ and *g*_*i*_ are not known, they will be assumed to be the sabe, that is, *f*_*i*_ = *f* and *g*_*i*_ = *g*. Eq. 6 themgreatly simplifies to the binomial of *n* = *n*_1_ + *n*_2_ +--- *n*_*N*_ with *k* = K,

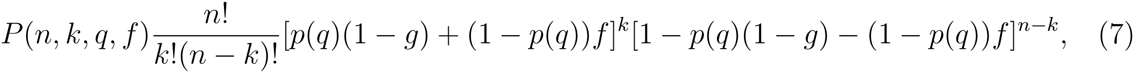

where now *n* = 462 and *k* = 41. The plot of this function for *f* = *g* = 0 can be found in Figure 5.

**FIG. 5:**
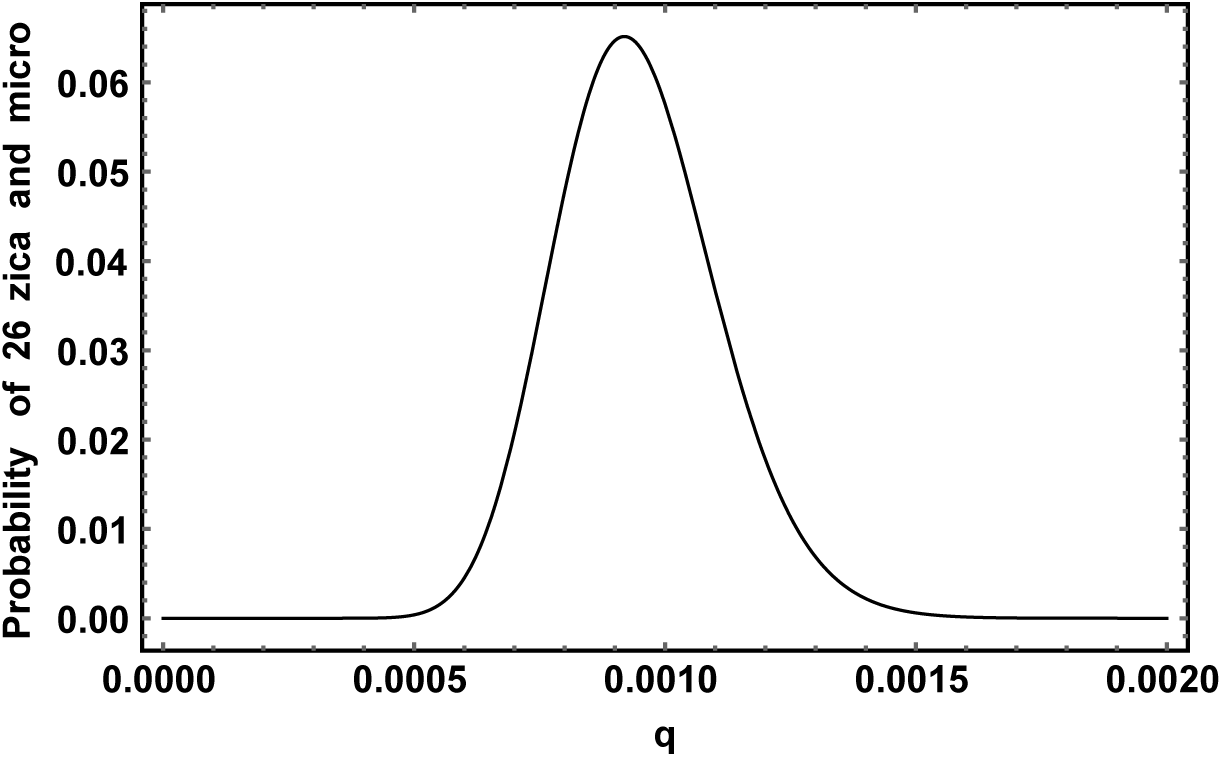
Probability of obtain 41 positives in a *n* = 462 sample of ZIKV and Microcephaly correlation study from ECDC.

Notice that this function is much more peaked and narrower than Figure 1, due to the fact it has more data points. From it one can get 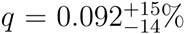. A search for the biggest probability as a function of *q*, *f* and *g* was done. Table II summarizes all results obtained.

This would suggest that the ZIKV causes Microcephaly in 0.12% of the pregnant women, the false positive to be only 0.6% but the false negative 84% of the cases. Again this numbers should be taken with care, as can be seen in Figure 6 where it was plot the P-Value as a function of a parameter t was used to relate *f* and *g* for illustrative purposes.

**FIG. 6:**
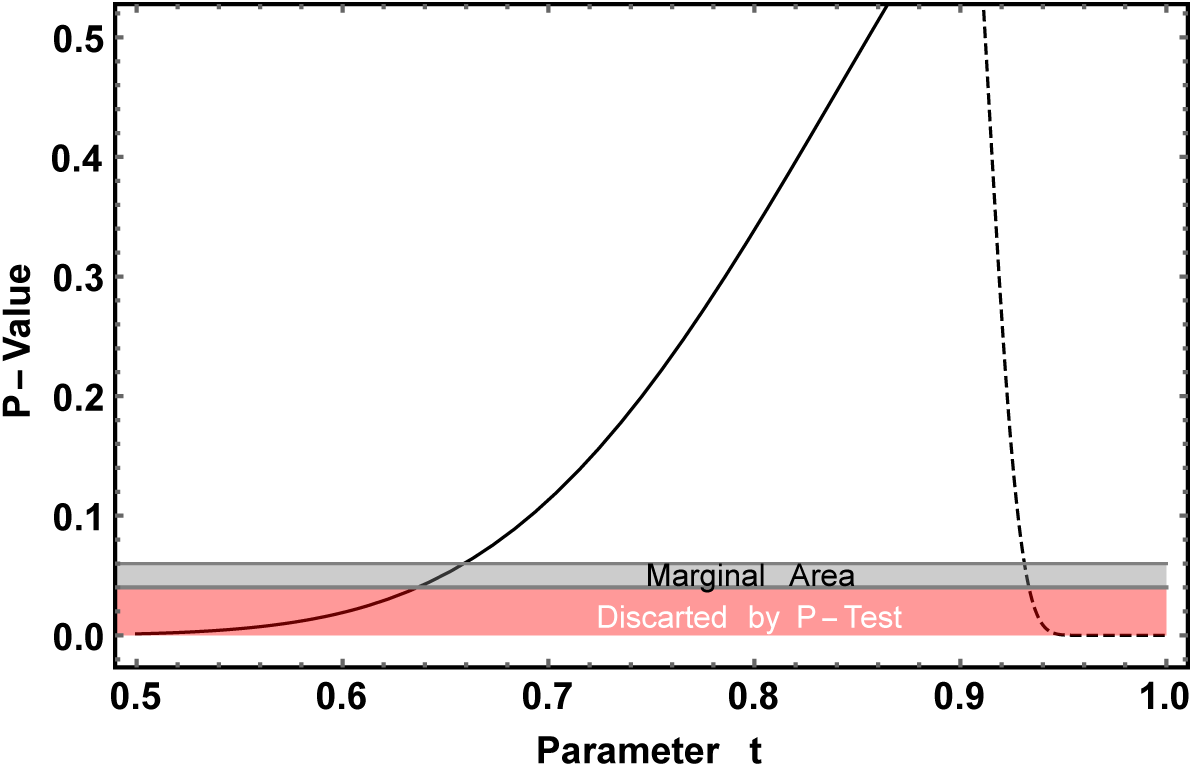
P-Value as a function of a parameter t. Full Line *q* = 0 and dashed line *q* = 0.00092.

Notice that it is possible to obtain a P-Value bigger than 0.05 even when *q* = 0, i.e. no correlation at all, even in this high statistic scenario. For example is enough that *f* = 6% and *g* = 30%, that is why it is so important to have a good measure of the reliability of the disease detection method.

## VI. CONSIDERATIONS ON FALSE POSITIVE TEST

Both results on the overall probability factor *q* = 2.3% and *q* = 0.09% make it that the methodology in each experiment (that is, *f* and *g*) has to be extremely clear and controlled to confirm a correlation between both diseases. Such correlation could already be asserted if those parameters were known. In practice they cannot be fully measured due to the fact that they involve individual genetic background, other infections, and environmental toxins, but at least an estimate considering the test methodology is fundamental.

An important point that has to be stressed out is the fact that even in case the correlation is shown to be true due to statistical analysis, it does not prove that there is a causal connection between them. A third factor could cause microcephaly and be connected to the ZIKV infection.

## VII. CONCLUSION

A statistical analysis was performed to probe the correlation between ZIKV and Microcephaly to check if it is possible to obtain relation between both. It was found that there seems to be a correlation between them where the probability of a pregnant woman with ZIKV to have a newborn with Microcephaly may be of 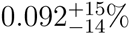. But even if such correlation exists, it is blurred by the ignorance of the false positive and false negative probability of tests for ZIKV in the mothers of infants. So futures tests of ZIKV should reach at least 30% of confidence to be able to distinguish those scenarios and a causal relation can only be asserted by viral inoculation tests.

